# *orco* mutagenesis causes loss of antennal lobe glomeruli and impaired social behavior in ants

**DOI:** 10.1101/112532

**Authors:** Waring Trible, Ni-Chen Chang, Benjamin J Matthews, Sean K McKenzie, Leonora Olivos-Cisneros, Peter R Oxley, Jonathan Saragosti, Daniel JC Kronauer

## Abstract

Life inside ant colonies is orchestrated with a diverse set of pheromones, but it is not clear how ants perceive these social cues. It has been proposed that pheromone perception in ants evolved via expansions in the numbers of odorant receptors (ORs) and antennal lobe glomeruli. Here we generate the first mutant lines in ants by disrupting *orco*, a gene required for the function of all ORs. We find that *orco* mutants exhibit severe deficiencies in social behavior and fitness, suggesting that they are unable to perceive pheromones. Surprisingly, unlike in *Drosophila melanogaster*, *orco* mutant ants also lack most of the approximately 500 antennal lobe glomeruli found in wild-types. These results illustrate that ORs are essential for ant social organization, and raise the possibility that, similar to mammals, receptor function is required for the development and/or maintenance of the highly complex olfactory processing areas in the ant brain.

---

Ants live in complex societies and display sophisticated social behavior, including reproductive division of labor between queens and workers, behavioral division of labor between nurses and foragers, the formation of adaptive foraging networks, nestmate vs. non-nestmate discrimination, and collective nest construction (David Morgan, 2009; Grüter & Keller, 2016; Richard & Hunt, 2013; Leonhardt et al., 2016). All of these behaviors are largely mediated via chemical communication using a wide range of pheromones (David Morgan, 2009; Grüter & Keller, 2016; Richard & Hunt, 2013; Leonhardt et al., 2016). However, the receptor families involved in pheromone perception have not been functionally characterized, in part because the complex life cycle of ants has hindered the development of functional genetic tools (Schulte et al., 2014; Grüter & Keller, 2016; Kohno et al., 2016; Reid & O'Brochta, 2016). In *Drosophila*, pheromone receptors have been identified that belong to multiple insect chemosensory receptor families, including odorant receptors (ORs), gustatory receptors (GRs), ionotropic receptors (IRs), and pickpocket channels (PPKs) (Kohl et al., 2015). Ants have numbers of GRs, IRs, and PPKs that are typical for insects, while their OR repertoire is highly expanded (Zube et al., 2008; C. D. Smith et al., 2011; C. R. Smith et al., 2011; Zhou et al., 2012; Oxley et al., 2014; McKenzie et al., 2016) (Figure 1A and Supplementary Table 1). This raises the possibility that the expansion of ORs specifically, rather than chemoreceptors in general, may underlie the evolution of complex chemical communication in ants. Ants also have exceedingly large numbers of glomeruli in their antennal lobes, which likely mirror their expanded OR gene repertoire (McKenzie et al., 2016) (Supplementary Table 1). Insect ORs function as chemosensory receptors by forming ligand-gated ion channels through the formation of dimers with the highly-conserved co-receptor protein Orco (Larsson et al., 2004; Jones et al., 2005; Sato et al., 2008). *orco* null mutants in fruit flies, locusts, mosquitoes, and moths therefore lose OR function and show impaired responses to odorants such as food volatiles and sex pheromones (Asahina et al., 2008; DeGennaro et al., 2013; Koutroumpa et al., 2016; Li et al., 2016; Yang et al., 2016).

**Figure 1:**
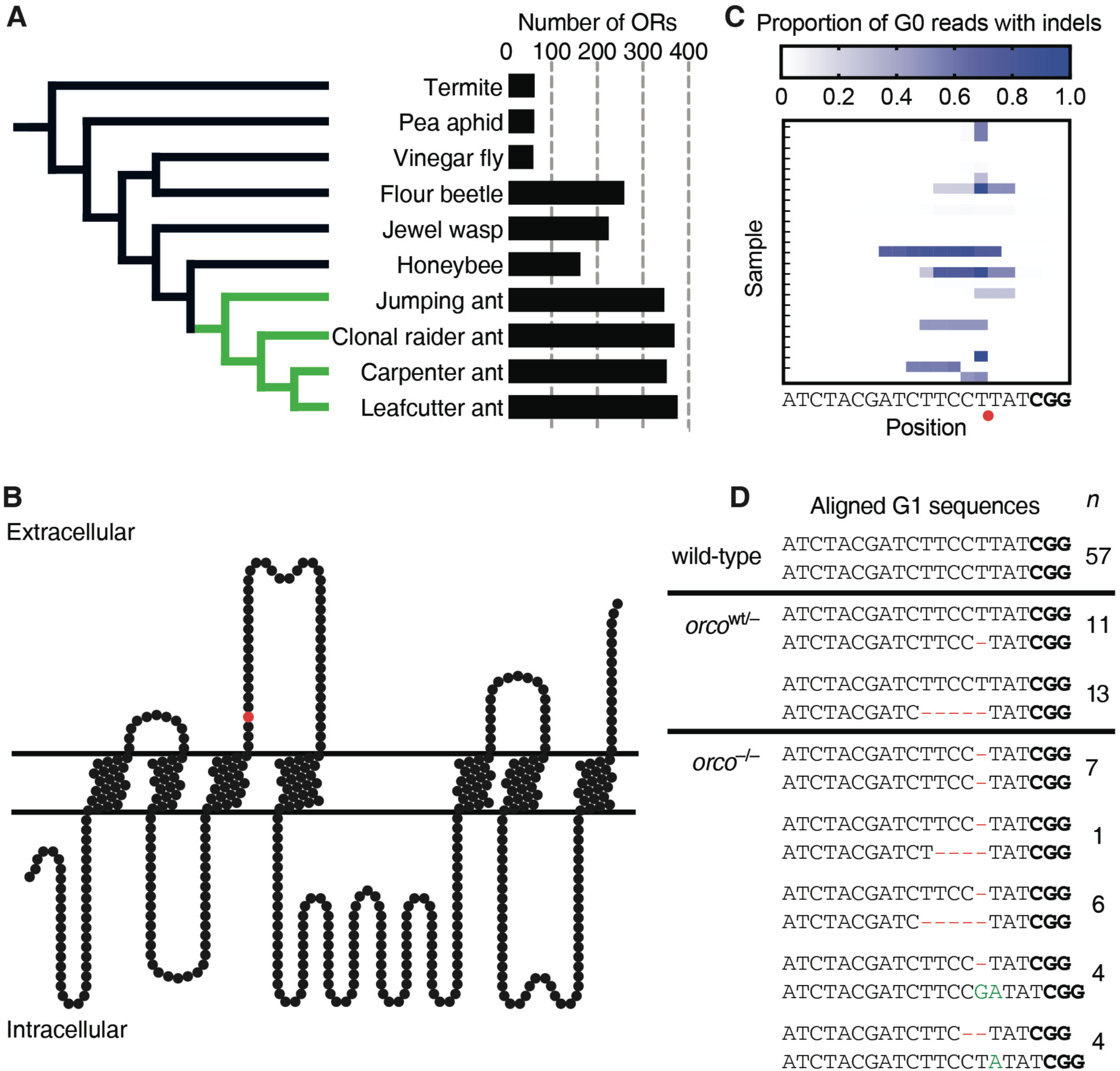
Number of OR genes and *orco* mutagenesis. (**A**) Phylogeny with numbers of ORs for ants (green) and other insects (black), showing ant OR expansion (Supplementary Table 1). (**B**) Position of predicted CRISPR/Cas9 cut site in Orco protein model (red circle). Frameshift mutations at this position truncate the wild-type protein between the third and fourth transmembrane domains, and the resultant mutant protein is unlikely to form functional ion channels. (**C**) Proportion of Illumina sequencing reads of *orco* amplicons with insertions or deletions (indels) relative to gRNA sequence in G0s, showing mutation rates of at least 97% in some individuals. Red circle indicates predicted CRISPR/Cas9 cut site. Protospacer adjacent motif (PAM) in bold. (**D**) Wild-type *orco* sequence compared to sequences for the two *orco*^wt/-^ and the five *orco*^-/-^ mutant lines based on G1 individuals. Deletions are shown in red and insertions in green. *orco*^-/-^ ants have two frameshift alleles and are therefore expected to be complete loss-of-function *orco* mutants. *n* indicates the number of ants of each line used across all experiments in this study. PAM in bold.

To study the role of ORs in ant biology, we developed a CRISPR/Cas9 protocol and created *orco* mutants in the clonal raider ant, *Ooceraea biroi* (formerly *Cerapachys biroi* (Borowiec, 2016)), a promising genetic model system (Oxley et al., 2014). We confirmed the identity of *orco* in the *O. biroi* genome (Supplementary Table 2, Supplementary Figure 1) and designed and synthesized a guide RNA (gRNA) to target *orco* (Figure 1B). We injected Cas9 and gRNA into 3,291 eggs less than 5h of age and produced 42 G0 adults, some of which displayed mutations in at least 97% of PCR amplicons of the *orco* target site (Figure 1C, Supplementary Text). G0 mutations in the germline can be inherited by G1s to produce stable modifications to the genome (Reid & O'Brochta, 2016). Given that *O. biroi* reproduces through parthenogenesis (Oxley et al., 2014), stable mutant lines can be clonally propagated from individual mutant G1s and subsequent generations without the need for crosses. *orco* loss-of-function mutant lines are thus derived from G1 eggs with independent frameshift mutations in both *orco* alleles. We recovered a diverse set of *orco* mutant lines, including two *orco*^wt/-^ lines with one frameshift allele and five *orco*^-/-^ lines with two frameshift alleles (Figure 1D). The phenotypes reported below were consistent across the two *orco*^wt/-^ lines and across the five *orco*^-/-^ lines, respectively. Descriptions of the specific lines used in each experiment along with the associated phenotypes are given in Supplementary Table 3.

In *D. melanogaster*, all antennal lobe glomeruli that have been examined remain present in *orco* mutants, implying that OR function is not required for gross antennal lobe morphology (Larsson et al., 2004; Chiang et al., 2009). These results contrast strongly with mice, where olfactory receptor function and neuronal activity are essential for the formation and maintenance of the analogous brain region, the olfactory bulb (Yu et al., 2004). However, *D. melanogaster* has only 60 ORs and a similar number of glomeruli, while mice possess over one thousand olfactory receptors and glomeruli. This striking difference suggests that highly complex olfactory systems must rely on receptor function for their development and/or maintenance. Ants have highly expanded numbers of ORs and antennal lobe glomeruli (Zube et al., 2008; Oxley et al., 2014) (Supplementary Table 1), raising the possibility that the development and/or maintenance of ant antennal lobes may require additional mechanisms to exceed the complexity found in other insects. To address whether OR function might be required for the structure of the ant antennal lobe, we imaged brains using confocal microscopy, measured antennal lobe volumes and the number of glomeruli, and reconstructed antennal lobes in wild-type and *orco* mutant adults in *O. biroi* and *D. melanogaster*. We found that the antennal lobes of *orco*^-/-^ ants measured only one third the volume of wild-type and *orco*^wt/-^ antennal lobes, and approximately 82% of the glomeruli were lost (Figure 2A,B and Supplementary Video 1). However, all six glomeruli in the T7 cluster of the ant antennal lobe, which is believed to be innervated by olfactory sensory neurons that do not express ORs (Nakanishi et al., 2010; McKenzie et al., 2016), were still present in *orco*^-/-^ individuals. No differences were observed in the volume of the protocerebrum, mushroom bodies, or central complex relative to wild-types (*P* = 0.45, 0.17, and 0.20, respectively; *t*-test). In contrast, we detected no significant difference in antennal lobe volumes and only minor potential differences in glomerulus numbers between wild-type and *orco*^-/-^ flies (Figure 2C,D, Supplementary Video 1, and Supplementary Text). These results demonstrate that development and/or maintenance of ant antennal lobes are indeed dependent on *orco* function.

**Figure 2:**
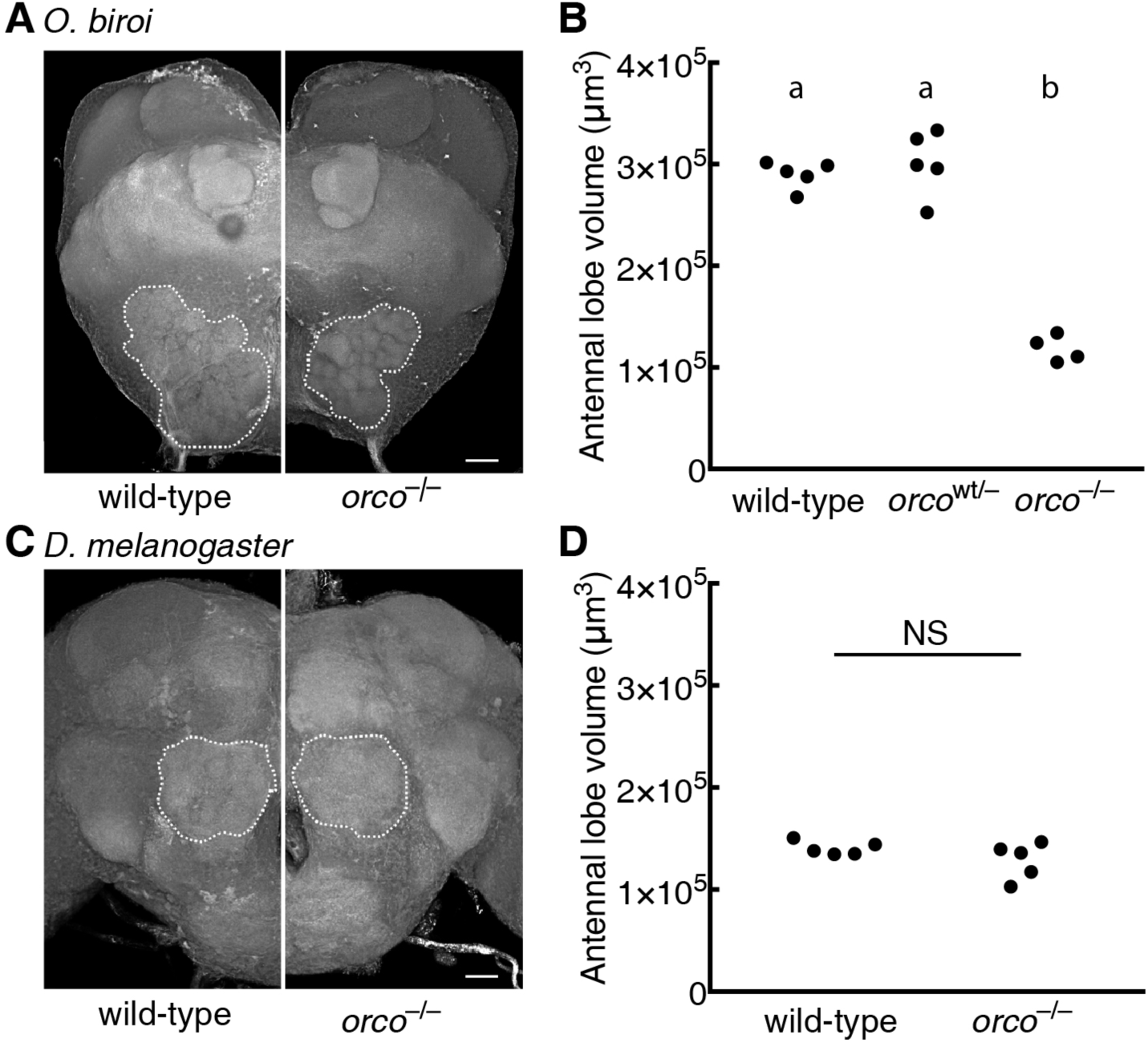
Reduced antennal lobes in *orco*^-/-^ ants. *O. biroi*: (**A**) Dorsal (n-ventral) 3D projections of wild-type and *orco*^-/-^ brains. Antennal lobes indicated by dashed lines. *orco*^-/-^ antennal lobe is highly reduced relative to wild-type. Two *orco*^-/-^ ants had 90 and 91 glomeruli relative to 493 and 509 glomeruli for two wild-type ants (one of the wild-type reconstructions has been published previously (McKenzie et al., 2016)); small differences between replicates within treatments might reflect reconstruction errors or actual biological variation. **(B**) Antennal lobe volumes for wild-type, *orco*^wt/-^, and *orco*^-/-^ ants. *orco*^-/-^ ants, but not *orco*^wt/-^ ants, have significantly smaller antennal lobes than wild-type. *D. melanogaster*: (**C**) Anterior (n-ventral) 3D projections for wild-type and *orco*^-/-^ brains. Antennal lobes indicated by dashed lines. *orco*^-/-^ antennal lobe is similar to wild-type. Two *orco*^-/-^ flies had 43 and 44 glomeruli, and two wild-type flies each had 46 glomeruli. These glomerulus numbers were higher than has been previously reported, which is likely due to differences in sample preparation and imaging techniques. Slight differences in glomerulus numbers between replicates may be due to reconstruction errors, or may reflect modest antennal lobe phenotypes in *orco* mutant flies (Supplementary Text). (**D**) Antennal lobe volumes for wild-type and *orco*^-/-^ flies. Volumes of wild-type and *orco*^-/-^ antennal lobes are not significantly different (*P* = 0.20, *t*-test). Scale bars are 20 μm. NS: not significant. Genotypic classes marked by different letters are significantly different (*P* < 0.05) after ANOVA followed by Tukey's test (B).

While further experiments are required to describe precisely how this striking phenotype arises, we hypothesize that ants, similar to mammals, may employ ORs in the development and/or maintenance of antennal lobe glomeruli, possibly allowing them to evolve larger and more complex antennal lobes than other insects (Supplementary Table 1).

Based on the general observation that ants are often repelled by the smell of permanent markers, we developed a simple assay to test whether *orco* mutants have compromised chemosensory abilities. We found that wild-type and *orco*^wt/-^ *O. biroi* are indeed strongly repelled by lines drawn with Sharpie^TM^ permanent marker, and rarely contact or cross Sharpie lines (Figure 3A-C). However, we found that *orco*^-/-^ *O. biroi* are significantly less repelled by Sharpie (Figure 3A-C and Supplementary Video 2), implying that *orco* is required to perceive the odorants that cause Sharpie lines to be repulsive. These results suggest that *orco* mutant ants possess general chemosensory deficiencies, similar to *orco* mutants in other types of insects (Asahina et al., 2008; DeGennaro et al., 2013; Koutroumpa et al., 2016; Li et al., 2016; Yang et al., 2016).

**Figure 3:**
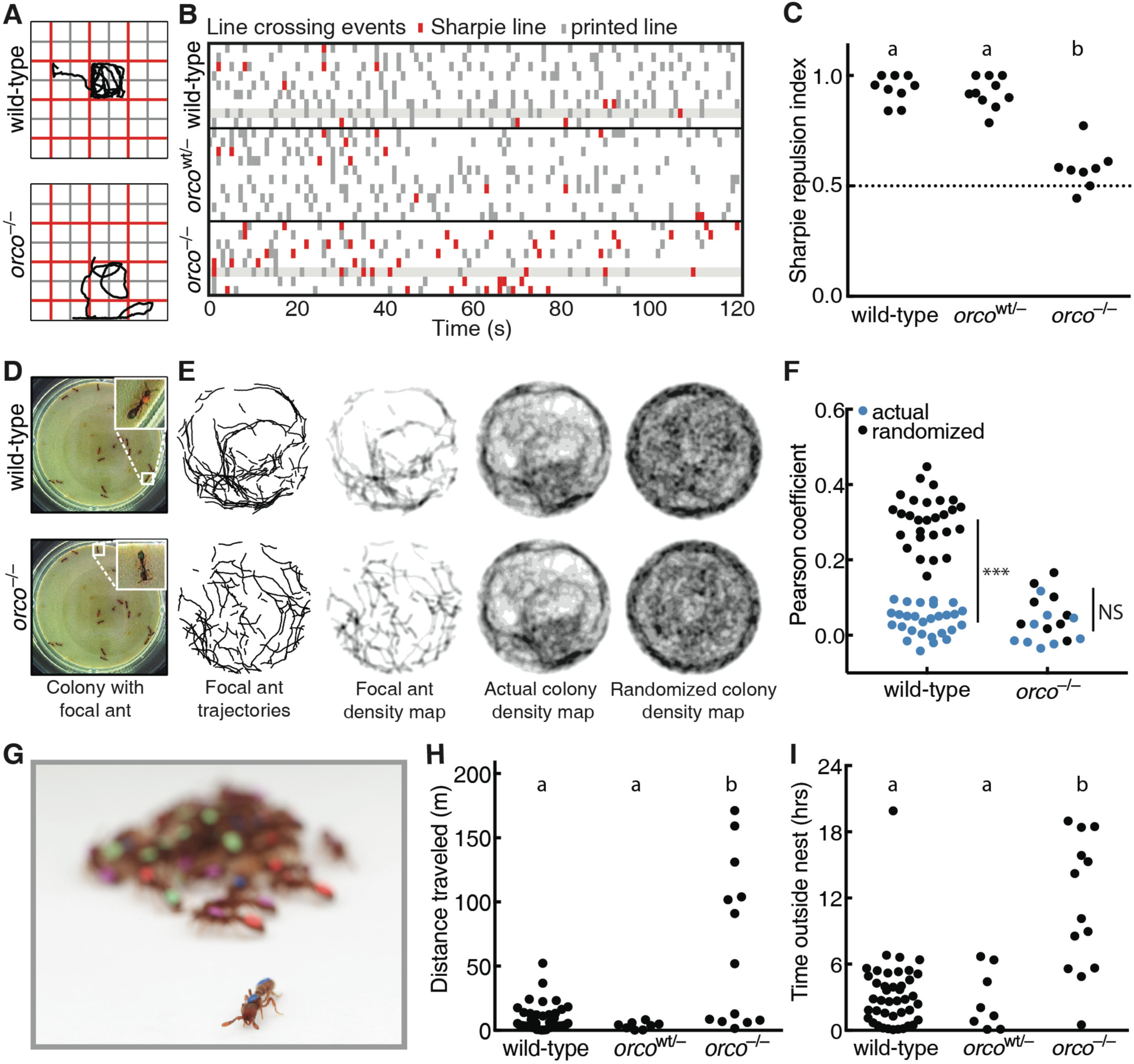
Deficient olfactory and social behavior in *orco*^-/-^ ants. (**A**) Example trajectories of wild-type and *orco*^-/-^ ants in Sharpie assay. (**B**) Line crossing events for wild-type, *orco*^wt/-^, and *orco*^-/-^ ants in Sharpie assays, with ants from Figure 3A highlighted. Wild-type and *orco*^wt/-^ ants cross Sharpie lines (red) less frequently than printed lines (grey), but *orco*^-/-^ ants cross both lines at approximately equal frequencies. (**C**) Repulsion indices for wild-type, *orco*^wt/-^, and *orco*^-/-^ ants in Sharpie assays. Repulsion index is calculated as proportion of printed line crosses. *orco*^-/-^ ants, but not *orco*^wt/-^ ants, are significantly less repelled than wild-types. (**D**) Example colony used for trail pheromone analysis. The same colony, containing a mixture of wild-type, *orco*^wt/-^, and *orco*^-/-^ ants, is shown twice, with a wild-type or *orco*^-/-^ focal ant highlighted. (**E**) Example trail pheromone analysis. Trajectories, during which ants were moving and edges were excluded, were used to create 2-D histograms, or density maps, for each ant in the colony. These density maps were compared to the actual and randomized density maps for all other ants in the colony. The wild-type density map is more strongly correlated with the actual colony density map than with the randomized colony density map, while the *orco*^-/-^ density map is poorly correlated with both colony density maps. (**F**) Pearson correlation coefficients for individual ant density maps with the actual or randomized density map of the other ants in the colony. Pearson correlation coefficients for wild-type ants, but not for *orco*^-/-^ ants, were significantly higher in actual than randomized density maps (sample sizes were too small to test *orco*^wt/-^). (**G**) Example colony showing an individual outside of the nest. (**H**) Distances traveled in 24 hr videos by ants in experimental colonies. *orco*^-/-^ ants, but not wild-type or *orco*^wt/-^ ants, exhibit a wandering phenotype. (**I**) Time without contacting other ants in 24 hr videos. *orco*^-/-^ ants spend more time without contact than wild-type or *orco*^wt/-^ ants. \*\*\**P* < 0.001; NS: not significant. Genotypic classes marked by different letters are significantly different (*P* < 0.05) after ANOVA followed by Tukey's test (C), or from log-likelihood ratio tests on generalized linear mixed models followed by Tukey's tests with colony as a random variable and actual/randomized maps (F) or genotypic class (H,I) as fixed variables.

Pheromone trails are a major feature of chemical communication in many ants, and are important for coordinating collective behaviors such as foraging and nest relocation (Zube et al., 2008; David Morgan, 2009). To test whether *orco* influences the ability of *O. biroi* to follow pheromone trails, we set up 5 experimental colonies composed of 12-14 identically-reared G1s with wild-type, *orco*^wt/-^, and *orco*^-/-^ genotypes, and individually tagged each ant with color dots (Figure 3D). We recorded videos of each colony and used a novel custom-built automated behavioral tracking system employing painted color tags (rather than paper barcodes (Mersch et al., 2013)) to individually identify the ants and quantify their behavior (Figure 3D, Supplementary Figure 2, and Supplementary Videos 3,4). We disturbed each colony at the beginning of each video. During the ensuing period of high activity, we created a 2-D histogram, or density map, of movement for each ant, and measured the Pearson correlation coefficient of this density map with the density map of the other ants in the colony, reasoning that density maps would be more highly correlated when ants were following pheromone trails (Figure 3E,F). To provide a null expectation, we also compared the density maps of individual ants to a randomized density map of other ants in the colony (Figure 3E,F). We found that the density maps of individual ants had significantly higher correlations with the density maps of the rest of the ants in the colony than with the randomized density maps in wild-type, but not *orco*^-/-^, ants (Figure 3F; average correlation coefficients were 0.31 versus 0.04 in wild-type and 0.07 versus 0.02 in *orco*^-/-^, respectively). These findings imply that trail following behavior is reduced or absent in *orco*^-/-^ ants, likely because they are unable to perceive chemical pheromone trails.

Nesting behavior, and the formation of aggregations more generally, is a ubiquitous feature of social insect biology (Depickere et al., 2004). Immediately upon eclosion, we noticed that some G1s did not nest with other ants, but instead showed a wandering phenotype (Figure 3G). In a set of 16 G1 colonies, we used this wandering phenotype to identify colonies containing *orco*^-/-^ ants with 100% accuracy (*P* < 0.001, Fisher exact test, see Methods). To more precisely measure nesting behavior in *orco* mutants, we recorded and analyzed 24 hr videos of each experimental colony. We found that wild-type and *orco*^wt/-^ ants aggregated into tight clusters and exhibited little movement outside the cluster, while *orco*^-/-^ ants frequently exited the cluster and wandered around the dish (Figure 3H, Supplementary Video 3). Overall, *orco*^-/-^ ants spent a significantly larger fraction of time without contact with other ants when compared to wild-type and *orco*^wt/-^ ants (Figure 3I). These findings demonstrate that typical nesting behavior is compromised in *orco*^-/-^ ants. This observation is consistent with the idea that *orco* mutants are unable to perceive odorants, such as aggregation pheromones (Bell et al., 1972; Depickere et al., 2004; Li et al., 2016), that might be involved in nesting behavior.

Finally, we investigated whether *orco* mutations have an effect on fitness by measuring egg-laying and survival rates of wild-type and *orco* mutant ants. We found that *orco*^-/-^ ants laid significantly fewer eggs than wild-types and *orco*^wt/-^ ants over a two week period (Figure 4A), and *orco*^-/-^ ants exhibited significantly higher mortality than wild-types over a 34 day period (Figure 4B). This suggests that the *orco* mutant phenotype has serious consequences for ant fitness (for a discussion of the potential role of off-target effects see Supplementary Text). It is possible that these fitness effects result because *orco*^-/-^ ants are unable to integrate into the colony, as wandering behavior and reduced fitness are also seen in wild-type ants that are kept in social isolation (Koto et al., 2015). While we observed many striking deficiencies in *orco*^-/-^ ants, it is important to note that these ants are viable, feed, lay eggs, and may still exhibit some typical social behaviors. For example, we have observed *orco*^-/-^ ants groom eggs, touch other ants with their antennae, and elicit alarm responses (Supplementary Videos 3,4). Thus, these *orco* mutants will provide an important resource to study the role of ORs in ant biology in the future.

**Figure 4:**
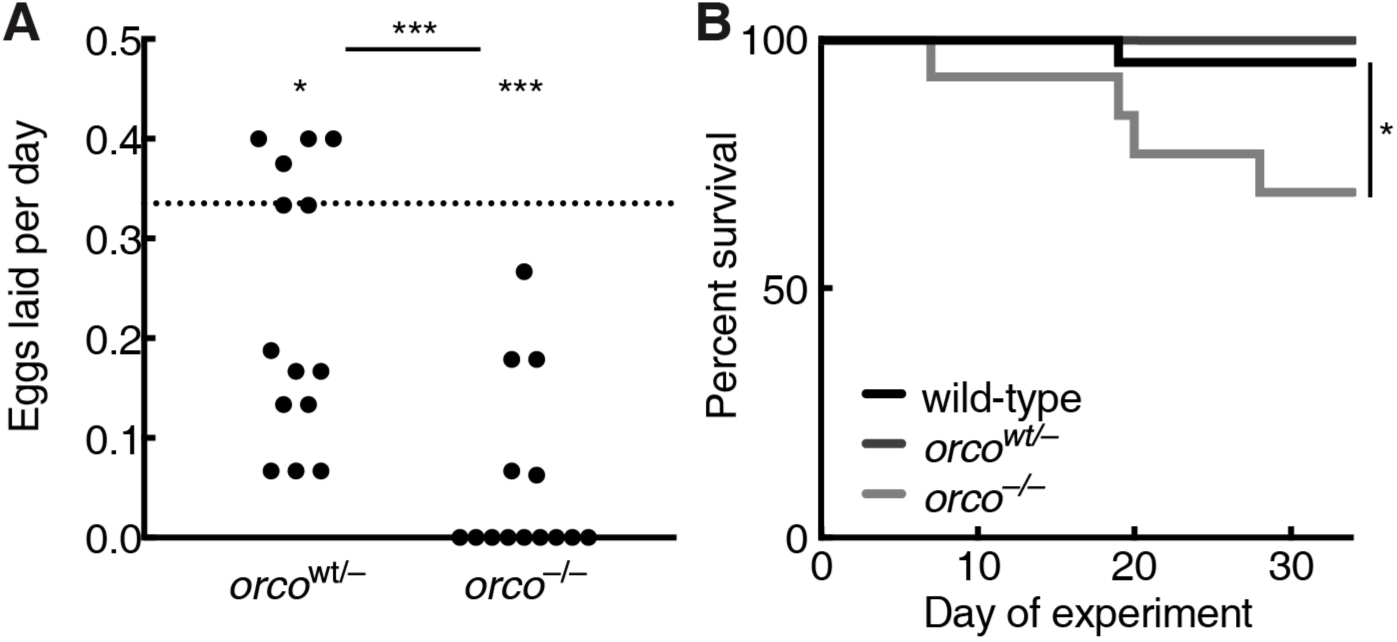
Reduced fitness in *orco*^-/-^ ants. (**A**) Eggs laid per day over a two week period by *orco*^wt/-^ and *orco*^-/-^ ants relative to wild-type average (dotted line). *orco*^-/-^ ants laid significantly fewer eggs than *orco*^wt/-^ ants. Both *orco*^wt/-^ and *orco*^-/-^ ants laid significantly fewer eggs than the wild-type average of 0.34 eggs per day. Wild-type data are given as an average, rather than individual values, because most ants in each colony were wild-type and it was therefore not possible to obtain individual egg-laying rates for wild-type ants (see Methods). (**B**) Survival of identically-reared and age-matched wild-type (*n* = 42), *orco*^wt/-^ (*n* = 7), and *orco*^-/-^ (*n* = 13) ants over a 34 day period. Survival of *orco*^-/-^ ants was significantly lower than that of wild-type ants. Survival of *orco*^wt/-^ ants was not statistically tested due to small sample size, but no trend toward reduced survival was observed. \**P* < 0.05; \*\*\**P* < 0.001; NS: not significant. *P* values from an unpaired two-way Wilcoxon test (comparison of *orco*^wt/-^ and *orco*^-/-^ egg-laying rates) and one-way Wilcoxon tests (comparisons of *orco*^wt/-^ and *orco*^-/-^ egg laying rates to wild-type) using the mean egg-laying rate of wild-type ants in this experiment (A) or from a Fisher exact test (B).

We have demonstrated that *orco* is crucial for many aspects of ant biology, including antennal lobe morphology, individual response to repulsive odorants and pheromones, and fitness. These results illustrate the functional significance of the striking expansion of ORs and antennal lobes in ants relative to other insects, and imply that the expansion of ORs may have been an important component of the evolution of eusocial behavior in ants (Zube et al., 2008; C. D. Smith et al., 2011; C. R. Smith et al., 2011; Zhou et al., 2012; Oxley et al., 2014; McKenzie et al., 2016) (Figure 1A, Supplementary Table 1). Major transitions in evolution require the coordinated action of individuals to operate as a functional, higher-level unit (Maynard Smith & Szathmary, 1997). During the transition from solitary to eusocial living in ants, this coordination was largely achieved via pheromones, and our study illustrates some of the dramatic consequences that are suffered by individuals deficient in pheromone perception. Importantly, the greatly reduced antennal lobes of *orco* mutant ants suggest that the mechanisms underlying the development and/or maintenance of ant olfactory systems differ from *D. melanogaster*, and could be dependent on receptor function as is the case in mammals. These findings also open the possibility that ants could serve as complementary models to better understand the evolution and development of complex chemosensory systems.

## Materials and Methods

### Confirmation of the *O. biroi orco* gene identity

Candidate *orco* orthologs for eight insect species were detected as reciprocal best hits using phmmer (Eddy, 1998) with *D. melanogaster orco* (flybase id: FBgn0037324) as the initial query sequence (Supplementary Table 2). To confirm orthology, homologs ±50% the length of *orco* were aligned with MAFFT (Katoh & Standley, 2013) using default parameters. This alignment was then used to construct a bootstrapped phylogeny with RAxML (Stamatakis, 2006), providing unambiguous support for a single copy ortholog of o*rco* in *O. biroi* (Supplementary Figure 1).

### Design of *orco* gRNA

Identification of cut sites and assessment of off-target sites was performed using the script cris.py, part of the genomepy package (commit #94cc628), available at https://github.com/oxpeter/genomepy. The genomic sequence for *O. biroi orco* was searched on both strands for the CRISPR guide RNA (gRNA) recognition sequence 5’ - N_20_NGG - 3’ using BLASTN (Altschul et al., 1990) and checked for off-target hits using CRISPRseek (Zhu et al., 2014). We detected no off-target sites with 2 or less mismatches and only one site with 3 mismatches for our *orco* gRNA, which is expected to lead to low or no off-target cutting (Fu et al., 2013) (see Supplementary Text).

### CRISPR reagent preparation

Recombinant Cas9 protein was purchased from PNAbio, and gRNA was synthesized as described previously (Kistler et al., 2015). Activity of Cas9 and gRNA was validated using an in-vitro digestion assay from New England Biolabs. Immediately prior to injection, Cas9 and gRNA were mixed in water to produce a solution with 100 ng/μL Cas9 and 10 ng/μL gRNA.

### Colony maintenance and egg collection

*O. biroi* colonies were maintained in circular Petri dishes (50 mm diameter, 9 mm height) with a plaster of Paris floor ca. 4 mm thick. Colonies were fed 3 times weekly with fire ant (*Solenopsis invicta*) brood and cleaned and watered at least once per week. Eggs for injections were collected from egg-laying units consisting of 70 *O. biroi* individuals without larvae or pupae. Eggs were collected and placed on double-sided tape with the ventral side up on a glass slide, and injected into the anterior end (Oxley et al., 2014). Slides were prepared with up to 80 eggs for injection and ~25 control eggs to validate incubation conditions. For additional information about egg collection see Supplementary Text.

### Egg injection, incubation, and rearing

Injection needles were prepared as in previous studies (Lobo et al., 2006). Injections were performed using an Eppendorf Femtojet with a Narishige micromanipulator. The Femtojet was typically set to Pi 1800 hPa and Pc 500 hPa. Needles were broken by gently touching the needle against a capillary submerged in halocarbon oil. Alternatively, sharper needles were generated by setting the Femtojet to maximum pressure (6000 hPa) and lightly touching the capillary against fibers on the tape. Data in this manuscript result from a combination of both methods.

Eggs were submerged in a drop of water immediately prior to injection, and dried after injection. Eggs were pierced with the needle and injected for 1-2 seconds. Eggs less than 5 hrs old were injected, corresponding to a time when eggs are in a syncytial stage of development with <100 nuclei (Oxley et al., 2014). We injected 100-300 eggs per day. After injection, eggs were incubated in humid plastic boxes prepared with a plaster of Paris floor moistened with distilled water.

Two *O. biroi* clonal lines, which are genetically distinguishable at the mitochondrial *cytochrome oxidase subunit 1* (CO1) gene (Kronauer et al., 2012), were used in this study. All experimental ants belonged to Line B, while Line A ants were only used as chaperones to raise experimental Line B individuals. Experimental ants were reared by placing Line B larvae (G0s) or eggs (G1s and subsequent generations) in colonies of 20 Line A chaperones, and chaperones were removed once the callows had eclosed. This rearing method results in a small fraction of Line A offspring of chaperones in colonies with the G0s and subsequent generations. For this reason, all individuals were genotyped following experiments, and Line A individuals were removed from all analyses. All experimental colonies in this study had eggs removed twice weekly so that adults were maintained without larvae or pupae. All individuals, including those that died during experiments, were genotyped (see below) to determine their clonal line, and *orco* amplicons from Line B individuals were sequenced to determine *orco* genotype. For additional information about egg injection, incubation, and rearing see Supplementary Text.

### Tagging

All ants in behavioral and fitness experiments (Figures 3, 4) were tagged with two color dots, one on the thorax and one on the gaster, using uni-Paint markers (models PX-20 or PX-21) such that each individual could be identified within the colony (Figure 3D,G). For automated behavioral tracking, four colors were used (blue, green, orange and pink) for a total of 16 unique combinations. Ants were tagged with a randomly assigned color pair at least 10 days prior to any behavioral experiments. Tagged ants had a leg removed for genotyping and sequencing either before (Figure 2A,B; Figure 3A-C) or after (Figure 3D-I; Figure 4) experiments.

### Genotyping

To distinguish Line A and Line B, eggs and adults were genotyped using PCR of mitochondrial *CO1* with standard DNA barcode primers (Folmer et al., 1994) followed by a restriction digest with MwoI from New England Biolabs. This enzyme cuts the PCR product derived from Line B, but not from Line A.

### Sequencing

To screen for *orco* mutations, we designed PCR primers that flanked the *orco* cut site and sequenced the resulting PCR products using Sanger and Illumina sequencing. Primer sequences were: F: <u>TCGTCGGCAGCGTCAGATGTGTATAAGAGACAG</u>TCCAACTTGCTGTAAATTTGGAT R: <u>GTCTCGTGGGCTCGGAGATGTGTATAAGAGACAG</u>CTCTTCTTGGTCGGCGGTA.

Illumina methods followed a previously described protocol (Kistler et al., 2015). Primers included tails at the 5’ end (underlined) that were used as adapters to add indices to individual samples for Illumina sequencing (Kistler et al., 2015). Sequences were aligned to the *orco* genomic sequence and reads at each base pair that aligned with an insertion or deletion were counted with the script crispralign.py from the genomepy package, available at https://github.com/oxpeter/genomepy. *orco* amplicons from 25 of 42 recovered G0s were subjected to Illumina sequencing (Figure 1C). Three of these individuals, all of which were found to have nearly 100% mutation rates, displayed a wandering phenotype similar to the wandering phenotype observed in G1s, indicating that somatic CRISPR in G0s may be useful for functional genetic studies even in the many social insect species where it is not logistically possible to generate or maintain stable mutant lines (Schulte et al., 2014).

### Identification of mutant sequences

Mutant lines were identified via Sanger sequencing of eggs and adults of G1s and subsequent generations. All Sanger sequencing traces were scored manually. For the G1 dataset (below), manual identifications were verified by checking for misalignment against a reference sequence using MEGA (Kumar et al., 2016) and by using the program Mutation Surveyor (Softgenetics) for automated allele identification. Mutant lines were defined as groups of ants that possess identical *orco* genotypes, and *orco* amplicons from representatives of each mutant line were Illumina sequenced and individual reads were manually inspected to ensure both alleles were properly identified.

### Glomerulus counts and antennal lobe volumes

One of the wild-type *O. biroi* antennal lobe reconstructions was based on published data (McKenzie et al., 2016). For the remaining data, *D. melanogaster* and *O. biroi* brains were dissected in PBS and immediately transferred to a fixative solution of either 1% glutaraldehyde or 2% PFA and 2.5% glutaraldehyde in PBS, and fixed at room temperature on a shaker for 1-30 days. To dehydrate, brains were rinsed in PBS and then suspended for 5 minutes each in an ascending series of 50%, 70%, 90%, 95%, 100%, 100%, and 100% ethanol. Brains were cleared and mounted in methyl salicylate. Glutaraldehyde-enhanced autofluorescence was imaged using a confocal laser scanning microscope (Zeiss LSM 8800) with excitation by a 488 nm laser. Three-dimensional projections were created from confocal image stacks using Fluorender (Wan et al., 2012). Three-dimensional reconstructions of glomeruli and antennal lobes were produced by manually segmenting confocal image stacks using the Segmentation Editor plugin in the Fiji distribution of ImageJ (Schindelin et al., 2012). Antennal lobe volumes were calculated using the Object Counter3D ImageJ plugin (Bolte & Cordelieres, 2006), blindly with respect to genotype.

### Sharpie assay

Sharpie assays were conducted with tagged ants (Figure 3A-C) on printer paper in a 5.25×5.25 in arena bounded by a clear acrylic barrier. Six horizontal and 6 vertical black lines were printed on the paper using an HP LaserJet printer (Figure 3A,B). Immediately before each assay, 3 alternating horizontal and vertical black lines were traced with red Fine Point Sharpie Permanent Marker (item 30002). Then the ant was placed in the center of the grid. A 2 min video was recorded and the number of times the ant crossed black and Sharpie lines was manually counted (Supplementary Video 2). A Sharpie repulsion index was calculated as the ratio of black line crosses to total line crosses. Once the experiment had concluded, we determined that low numbers of line crosses caused the repulsion indices to be unreliable, and we therefore excluded four assays that had less than 10 line crosses total. As a positive control, a wild-type worker was assayed after each assay with a low repulsion index to ensure the Sharpie lines retained a repulsive effect. All positive controls had high repulsion indices and as a population were statistically indistinguishable from the other wild-type workers assayed (*P* = 0.42, *t*-test).

### G1 preparation for behavior and fitness experiments

G1 rearing resulted in a set of 34 colonies containing a mixture of G1 ants and Line A progeny of chaperones. These colonies were used to identify mutants for egg-laying, automated behavioral tracking, and survival experiments (Figure 3D-I, Figure 4, Supplementary Table 3). Once each colony started producing eggs, we collected all eggs 5 times over a 14-16 day period. *CO1* amplicons from all eggs were genotyped to identify Line A and Line B eggs, and *orco* amplicons from Line B eggs were Sanger sequenced to identify *orco* mutants. We sequenced 2,184 eggs from the 533 Line B ants in these colonies, corresponding to ca. 4 eggs per ant. During the period of egg collection and one week after egg collection had concluded, we subjectively determined whether any individuals in any given colony displayed a wandering phenotype. Colonies in which *orco* mutant eggs were detected or in which wandering phenotypes were observed were selected for the egg-laying dataset.

### Egg-laying dataset

We included 16 colonies in the egg-laying dataset. A subset of ants in these colonies were later also used for behavioral and survival experiments (Supplementary Table 3, see below). After the experiments had been concluded, all ants had a leg removed, from which *orco* amplicons were sequenced. For each *orco*^wt/-^ and *orco*^-/-^ adult we identified, we counted the number of eggs of its genotype in its colony. If several adults of the same genotype were identified in a colony, for each individual we calculated the number of eggs of that genotype divided by the number of adults of that genotype. We used a two-tailed Wilcoxon test to test whether *orco*^wt/-^ or *orco*^-/-^ G1s produced different numbers of eggs than the average of wild-type G1s in this experiment (Figure 4A).

### Behavioral and survival dataset

Before removing legs from ants for genotyping, workers from colonies that produced a high frequency of *orco* mutant eggs or contained individuals with wandering phenotypes were pooled to create 5 experimental colonies with a mixture of 12-14 G1 wild-type and *orco* mutant ants. Before pooling, all workers in these colonies were individually tagged with two color dots. These 5 colonies were recorded in 24 hr videos.

Experimental *O. biroi* colonies initially contained a total of 68 G1 ants, with 42 wild-type, 8 *orco*^wt/-^, and 14 *orco*^-/-^ individuals (Figure 1D). These colonies also contained 4 *orco* mutant individuals with in-frame mutations, which were not included in the current analyses because sample sizes were too small. G1s in experimental colonies varied in *orco* genotype but were otherwise identical in rearing methods, genetic background, and did not differ systematically in age. Before the start of each 24 hr video, colonies were cleaned and the plaster was moistened. For four weeks after the video was recorded, we also recorded survival of all ants in the 5 experimental colonies.

### Video recording and automated behavioral tracking

Automated behavioral tracking was performed in custom-made tracking setups under constant illumination. Temperature was maintained at 25°C. Videos were acquired using C910 Logitech USB webcams controlled with custom MATLAB (version R2016a, The MathWorks, Inc.) software at 10 fps at 960×720 pixel resolution (13 pixels/mm).

Tracking was performed blind with respect to genotype. Videos were processed and analyzed using custom MATLAB software. In each frame, ants were segmented from the background of the dish using a fixed threshold. Resulting components, or blobs, were linked into trajectories using the optical flow computed between consecutive frames (Horn-Schunck method (Barron et al., 1994)). Trajectories ended and new ones were initiated whenever blobs split or merged between two consecutive frames. Trajectories stored the following data, collected from the respective blob in each frame: centroid (position of center of mass), orientation (angle between the major axis of the best-fitting ellipse and the horizontal axis) and area (in mm^2^).

We used a threshold size to select trajectories that corresponded to a single ant and lasted longer than two seconds. Each trajectory was then assigned a combination of color tags using a custom classification algorithm. For each experiment, at least 500 frames per ant were manually identified to create a training set, 70% of which was used to train, and 30% of which was used to validate an automated identity classifier.

For each trajectory, a naive Bayes color classifier (Fletcher et al., 2011) was used to compute the pixel color probabilities for each pixel in the blob of each frame for six color classes: the four tag colors, the ant cuticle color and the color of the plaster of Paris (Supplementary Figure 2). Predicted probabilities for all four tag colors were used to determine whether both tags were visible and, if so, the orientation of the ant in the frame was deduced from the relative position of the tags with respect with the cuticle color. If both procedures were successful, the pixel color probabilities were fed into another naive Bayes classifier to assign an identity to the ant.

For each trajectory, frames were tested in a random order until 20 frames were successfully identified or no more frames were available. If at least one frame was identified, the trajectory, and thus all its underlying positions, was assigned the most frequently predicted identity. The identity classifier had an empirical error rate of 20% on single frames. However, the error rate decreased with the number of frames tested within a trajectory. Overall, we estimate that less than 10% of the total identified positions were misclassified, a performance equivalent to that reported previously for a functionally similar tracking setup for ants (Mersch et al., 2013).

### Behavioral analysis

Cleaning and watering the nests at the beginning of each experiment caused the ants to actively move around the dish. We used this initial period of high activity to measure trail following behavior, reasoning that trail pheromones may cause two or more ants to move along the same path within the dish. For each ant, we measured the correlation between its own movement and the movement of the remaining ants in the colony during the first hour of the tracking experiment. As *O. biroi* have a tendency to walk along the edges of the dish, we discarded segments of trajectories close to the edge of the dish and included only segments where the ants moved faster than 1 mm/s continuously for at least 1 second and without contact with any other ant. We then computed a 2-D histogram, or density map, for each ant by counting the number of times one of the remaining positions in the 970×720 pixel original image fell into each bin of a 120×90 bin grid. We then computed the Pearson correlation coefficient between the density map of each ant and a density map constructed from the trajectories of all ants in the colony but the focal ant (Figure 3E). For each ant, the actual correlation coefficient provided an estimate of the correlation of movement of that ant with the actual movement of other ants in the colony, which presumably results from following pheromone trails.

As a baseline comparison, the density map of each ant was correlated with a randomized density map constructed by rotating the trajectories of all ants but the focal ant in the colony by a random angle around the center of the dish (Figure 3E). This randomized correlation coefficient provided an estimate of the correlation of the movement of that ant with the randomized movement of other ants in the colony. This residual correlation reflects the portion of the correlation that is due to non-local effects such as turning frequencies, linear and angular velocity dynamics, and radial preference for certain regions of the petri dish. After examining the data, two experimental colonies were excluded from the trail following analysis because they did not form clear trails during the videos, resulting in Pearson correlation coefficients of approximately zero for all ants in the colony.

To measure wandering phenotypes, in each experiment we calculated the total distance traveled by each ant over the 24 hr video by computing the distances between all pairs of successive positions in meters in all identified trajectories. Time without contact was calculated as the total time each ant was identified in each experiment (since ants were only identifiable when they were separated from other ants). This provides a minimum estimate of the time without contact for each ant, given that it was also possible for ants to be spatially separated from other ants, yet unidentifiable, for example if their posture did not allow the detection of both tags.

### Statistics

Behavioral tracking and antennal lobe volume measurements were performed blindly with respect to genotype. Other analyses were not performed blindly with respect to genotype. Mixed model statistics were performed in R v 3.3.1 using the *lmer* function in the lme4 library as described previously (Ulrich & Schmid-Hempel, 2012; Ulrich & Schmid-Hempel, 2015). All other statistics were performed using GraphPad Prism 7. Normality was determined by D'Agostino-Pearson normality tests. Datasets used for ANOVA analyses had equal variance across treatments. Single groups were compared against a predicted mean using two-tailed Wilcoxon tests. Proportional data were compared between treatments using a Fisher exact test. Unpaired two-tailed Student's *t*-tests or paired Wilcoxon tests were used to compare two groups, when appropriate, and two-way ANOVAs followed by Tukey's multiple comparisons test were used to compare more than two groups.

### Data and code availability

Raw data are available in Supplementary Table 3. gRNA design and sequencing analyses were performed using the script crispralign.py from the genomepy package, available at https://github.com/oxpeter/genomepy.

## Acknowledgments

We thank the members of the Kronauer, Vosshall, and Ruta laboratories for reagents and critical insights. We are grateful to Ingrid Fetter-Pruneda for suggesting the Sharpie experiment, Yuko Ulrich for help with mixed-model statistics, Laura Seeholzer and Margaret Ebert, along with the Insect Genetic Technologies Research Coordination Network, for help in developing the injection protocol, Vanessa Ruta for providing flies, Leslie Vosshall for access to laboratory equipment, Zak Frentz and Asaf Gal for help in implementing the automated behavioral tracking, as well as Cori Bargmann, Claude Desplan, and Leslie Vosshall for comments on a previous version of this manuscript. This work was supported by grant 1DP2GM105454-01 from the NIH, a Searle Scholar Award, a Klingenstein-Simons Fellowship Award in the Neurosciences, and a Pew Biomedical Scholar Award to D.J.C.K. P.R.O. was supported by a Leon Levy Neuroscience Fellowship, and J.S. and S.K.M. were supported by a Kravis Fellowship and NRSA Training Grant #GM066699, respectively. W.T. and D.J.C.K. conceived the project with input from B.M. W.T., B.M., and L.O.C. conducted preliminary experiments to develop CRISPR methods in *O. biroi*. W.T., N.C., and L.O.C. performed injections and ant rearing. W.T., N.C., L.O.C., and P.R.O. performed sequencing and analysis with input from B.M. L.O.C. performed brain dissections, and S.K.M. performed remaining neuroanatomical experiments and analyses. W.T., L.O.C., N.C., and J.S. performed behavioral experiments. J.S. developed automated tracking techniques and performed behavioral analyses involving automated tracking. W.T., N.C., and L.O.C. performed fitness experiments. D.J.C.K. supervised the project. W.T. and D.J.C.K. wrote the paper. All authors discussed the results and the manuscript. This is Clonal Raider Ant Project paper #6.

## Supplementary Text

### *D. melanogaster* glomerulus counts

Our reconstructions of *D. melanogaster* antennal lobe glomeruli yielded different glomerulus numbers than what has been published previously (Supplementary Table 1). Our reconstructions also showed small differences in glomerulus numbers between wild-type and *orco*^-/-^ flies (Figure 2). To address this possibility, we imaged and reconstructed two additional *D. melanogaster* antennal lobes, one from an *orco*^-/-^ and one from a wild-type individual. These reconstructions were not performed strictly *de novo*, as the *O. biroi* and *D. melanogaster* reconstructions reported in the main text, but by referring to the published map of the *D. melanogaster* antennal lobe (Laissue et al., 1999; Laissue & Vosshall, 2008). Due to differences in sample preparation and imaging methods, it was not possible to unambiguously match each glomerulus to the published map. However, we identified structures in our newly reconstructed wild-type and *orco*^-/-^ antennal lobes that corresponded to all published wild-type glomeruli. These results suggest that *orco*^-/-^ flies have no systematic reduction in the number of antennal lobe glomeruli compared to wild-types, although it is possible that some neighboring glomeruli in *orco*^-/-^ flies have divisions that appear less distinct or may even be fused relative to wild-types (Figure 2).

### Off-target effects

It has been shown that in some cases CRISPR/Cas9 injections can lead to off-target mutagenesis that in turn can give rise to non-specific phenotypes. However, we consider this to be highly unlikely in the current study due to the following reasons. First, we used a high-quality reference genome to design the gRNA in this study to have no additional target sites in the genome that are likely to lead to off-target cutting (Fu et al., 2013; Oxley et al., 2014) (see Methods). Second, mutations induced by Cas9 are stochastically generated (Fu et al., 2013), such that any off-target effects would likely be present in some G1 lines but not others. The phenotypes we report are consistent across five independently generated *orco*^-/-^ lines, and we do not observe the same phenotypes across two independently generated *orco*^wt/-^ lines (Supplementary Table 3). Third, the striking reduction of *orco*^-/-^ antennal lobes relative to *orco*^wt/-^ and wild-type antennal lobes is a phenotype specific to the ant chemosensory system that is unlikely to arise from random off-target effects. This phenotype provides a direct functional link between the *orco*^-/-^ genotype and the chemosensory deficiencies described in this study. Importantly, the antennal lobe phenotype was entirely discrete: every *orco*^-/-^ brain had substantially smaller antennal lobes than any *orco*^wt/-^ or wild-type brain, even though this phenotype was measured across multiple independently derived *orco*^-/-^ and *orco*^wt/-^ lines (Figure 2b, Supplementary Table 3). Therefore, while we cannot exclude the possibility that our injections gave rise to some level of off-target mutations, it is unlikely that the specific phenotypes reported in this study arise from off-target effects.

### Additional egg collection, injection, and rearing methods

We observed that the presence of eggs in *O. biroi* colonies inhibits the production of new eggs and employed this observation for efficient egg collection. Egg-laying units were left with eggs for 7 days to inhibit worker egg-laying. On day 0 eggs were removed to release inhibition, and on day 2 eggs were removed again to further prevent inhibition. This led workers to synchronously activate their ovaries, and on days 5, 6, and 7 eggs were collected for injection. Following day 7, eggs were not collected from these colonies for 7 days, causing workers to become inhibited, and the protocol was then repeated.

Eggs were collected from colonies under a stereoscope using insect pinning needles. On a typical injection day, eggs were removed from colonies from 10-11 am, and those eggs were used as uninjected incubation controls or fostered into rearing units. Eggs were collected for injections from 2-3 pm and 6-7 pm and injected from 3-4 pm and 7-8 pm, respectively. Therefore, all injected eggs were less than 5 hrs old, when *O. biroi* eggs are in a syncytial stage of development with ?100 nuclei (Oxley et al., 2014). Typical egg-laying units produced 2-5 eggs per day, and we collected from up to 60 egg-laying units, injecting 100-300 eggs per day.

The anterior end of *O. biroi* eggs is slightly wider than the posterior end, and the ventral surface is concave while the dorsal surface is convex. To inject, eggs were placed on double-sided 3M tape (Model S-10079 from Uline) on a glass slide, with the anterior end forward and the ventral side upward. Eggs were injected into the anterior end, where nuclei are located in early *O. biroi* embryos (Oxley et al., 2014). The ventral side was placed upward, so that larvae hatched with the mouth facing away from the tape, which facilitated successfully recovering larvae from the tape. To inject, eggs were individually submerged in 1-2 μL drops of water. Eggs were gently pierced with the needle and injected for 1-2 seconds. During successful injections, little or no cytoplasm is discharged from the egg when the needle is removed. Preliminary trials showed that injection under liquid was necessary to remove the needle without rupturing the chorion, and that water led to higher survival than Ringer, PBS, or halocarbon oil. A video was recorded of every injection session, allowing us to verify that hatching larvae had been successfully injected.

Preliminary injections were conducted using multiple batches of reagents and variable Cas9 and gRNA concentrations. These trials suggested that hatch rates varied inversely with cutrates. Batch effects in hatch rates were observed across different days of injection within the same experimental treatment, requiring multiple injection days and large numbers of eggs (~400) to accurately estimate hatch rates of any given experimental treatment. In the final injection round, eggs were injected with either low (1800 hPa) or high (6000 hPa) pressure (see Methods), with sharper needles and lower injection volumes used in injections with high pressure. 46 of 2535 eggs (1.8%) hatched after injections with low pressure, and 58 of 756 eggs (7.6%) hatched after injections with high pressure. 25 of the 42 G0s were Illumina sequenced, and we observed an average of at least 27% cutrates at the predicted cut site resulted from low pressure injections (*n* = 17) relative to 22% from high pressure injections (*n* = 8). *orco*^wt/-^ and *orco*^-/-^ G1s were recovered from G0s injected with each method.

Following injections, slides with eggs were incubated in air-tight plastic boxes (0.9L SpaceCube boxes from ClickClack). Incubation boxes were prepared with a plaster of Paris floor (85 g plaster of Paris mixed with 50 mL distilled water). The plaster was dried completely after casting, and water was then added until the plaster became saturated with moisture to determine the saturation volume. The plaster was then dried completely once again, after which 20% of the saturation volume of distilled water was added. This procedure produced suitable incubation conditions for 2 weeks, after which the plaster was discarded. Incubating eggs were checked daily, and any water that had condensed on the eggs was removed with Kimwipes^TM^ tissue. Fungus frequently grew on injection slides. Growth was controlled by spacing the eggs ~2 mm apart and mechanically breaking up fungal hyphae and overgrown eggs in 100% ethanol using insect pinning needles. This egg-incubation protocol yielded ~60% hatch rates of uninjected control eggs, which is similar to hatch rates of eggs in laboratory colonies.

To synchronize hatching of larvae from injected eggs, eggs injected on days 5, 6, and 7 were incubated at different temperatures. Preliminary trials showed that eggs incubated at 25 °C hatch after 9-10 days, while eggs incubated at 30 °C hatch after 7-8 days. We therefore incubated eggs injected on days 5 and 7 at 25 °C and 30 °C, respectively, while eggs injected on day 6 were incubated at 25 °C for the first 5 days and then at 30 °C until hatching. This protocol resulted in most larvae hatching on days 14 and 15. Once hatching had commenced, larvae were manually removed from the egg membrane with an insect pinning needle, taking care to prevent them from becoming stuck to the double-sided tape. Eggs that were expected to hatch overnight were wrapped with a sheet of Parafilm^®^ (stretched to be as thin as possible) to prevent larvae from falling onto the tape.

To rear G0 larvae, uninjected control eggs slightly older than the eggs injected on day 5 were placed with ~20 adult Line A workers in a Petri dish with a plaster of Paris floor and maintained at 25 °C. These eggs hatched slightly earlier than the injected eggs, priming the workers to rear larvae derived from injected eggs. When the larvae hatched from injected eggs, control larvae were replaced with experimental (injected) larvae. Preliminary trials showed that higher survival was obtained by fostering a minimum of 7 larvae at a time, so control larvae were added to experimental larvae if insufficient experimental larvae were available. The G0 adults reported in this study therefore include an unknown fraction of adults derived from control larvae. Survival of larvae under these conditions was approximately 50%.

---

**Supplementary Figure 1:**
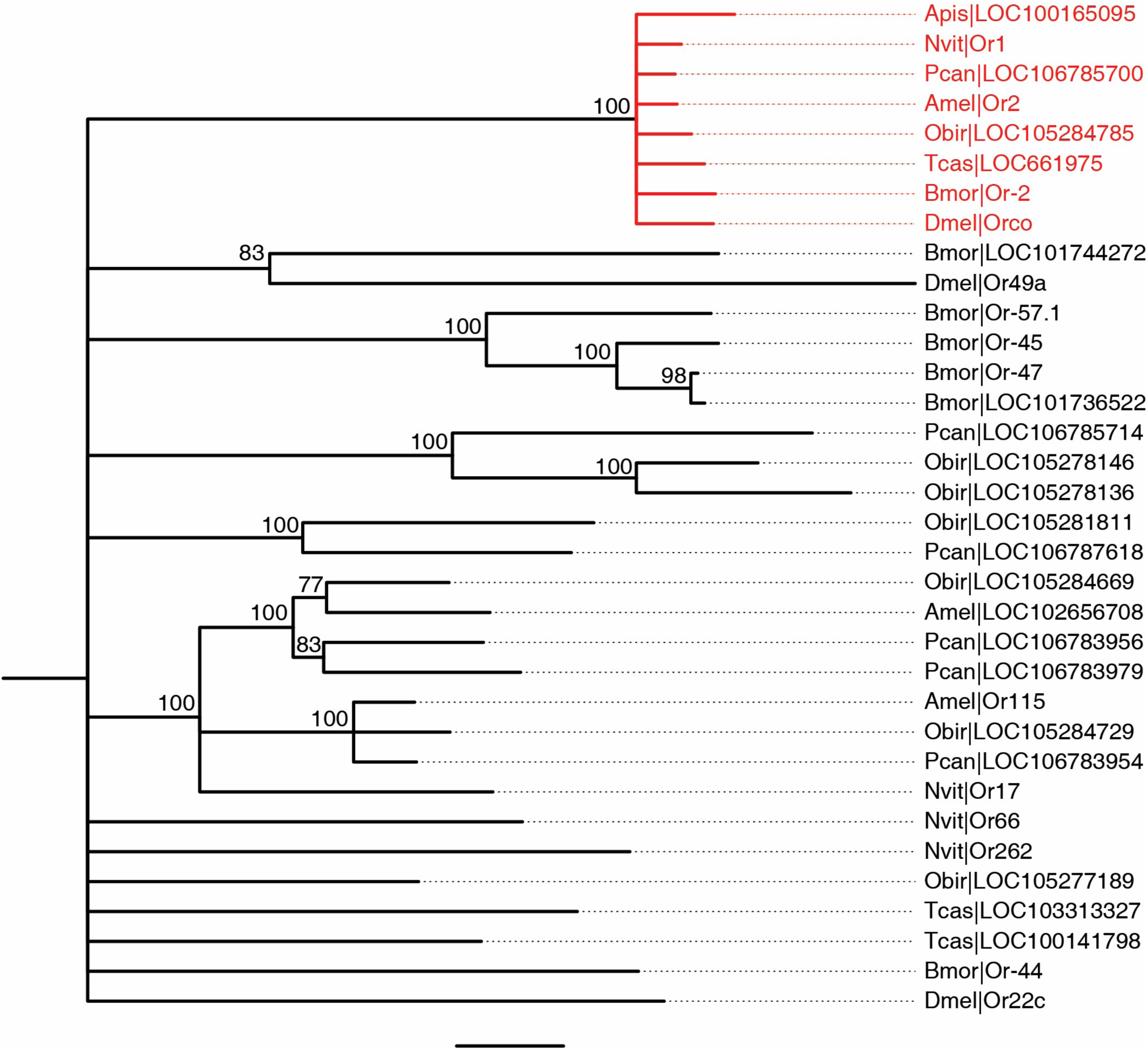
Phylogeny of *orco* and other odorant receptor genes in insects. *orco* orthologs in red. *O. biroi orco* is included in a clade with the *orco* genes from all other studied insect species (100% bootstrap support). All nodes with less than 75% bootstrap support have been collapsed for clarity. All other bootstrap values are shown at the respective node. Species are indicated by a four letter code: Apis - *Acyrthosiphon pisum*; Nvit - *Nasonia vitripennis*; Pcan - *Papilio canadensis*; Amel - *Apis mellifera*; Obir - *Ooceraea biroi*; Tcas - *Tribolium castaneum*; Bmor - *Bombyx mori*; Dmel - *Drosophila melanogaster*. NCBI gene symbols are shown on the right. The scale bar indicates an average of 0.3 substitutions per site.

**Supplementary Figure 2:**
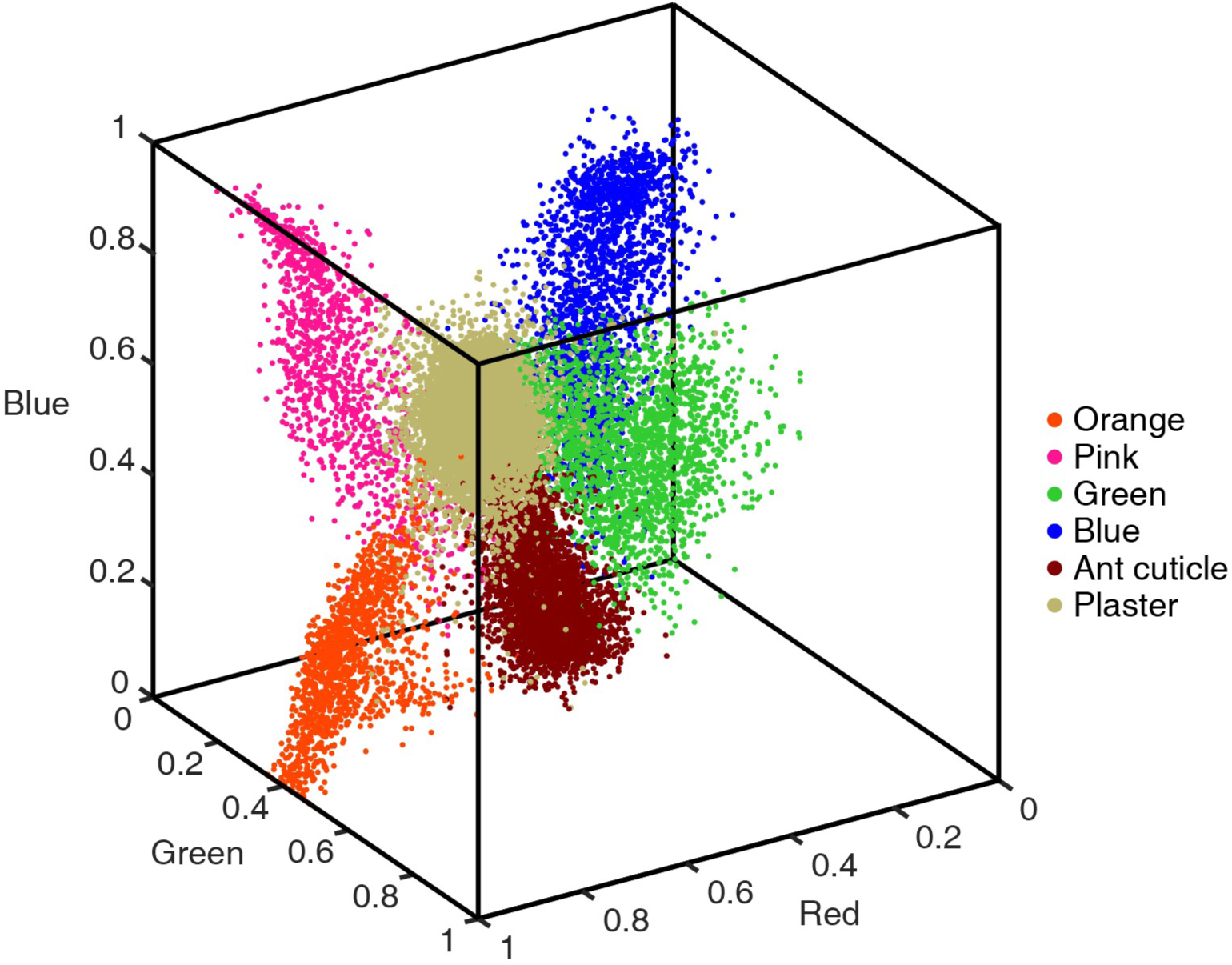
Color definition for automated behavioral tracking in RGB space. Measurements were collected manually from the four different tag colors (orange, pink, green and blue), the plaster (background) and the ant cuticle. These colors are clearly distinguishable in RGB space, allowing automated behavioral tracking based on identifying the color tags of each ant.

**Supplementary Table 1:** OR and glomerulus numbers for ants and other insects. For social insects, glomerulus counts refer to workers. OR numbers refer to putatively functional genes.

| Common name | Scientific name | OR number | Antennal lobe glomerulus number |
| --- | --- | --- | --- |
| Dampwood termite | *Zootermopsis nevadensis* | 63 (Terrapon et al., 2014) | 70-74 (Terrapon et al., 2014) |
| Pea aphid | *Acyrtosiphon pisum* | 62 (Smadja et al., 2009) | 25-40 (Kollmann et al., 2011) |
| Vinegar fly | *Drosophila melanogaster* | 60 (Robertson et al., 2003) | 42 (Laissue et al., 1999; Laissue & Vosshall, 2008) |
| Red flour beetle | *Tribolium castaneum* | 259 (Engsontia et al., 2008) | ~70 (Dreyer et al., 2010) |
| Jewel wasp | *Nasonia vitripennis* | 225 (Robertson et al., 2010) | Not published |
| Honey bee | *Apis mellifera* | 163 (Robertson et al., 2010) | 156-166 (Flanagan & Mercer, 1989) |
| Jerdon's jumping ant | *Harpegnathos saltator* | 347 (Zhou et al., 2012) | 178* (Hoyer et al., 2005) |
| Clonal raider ant | *Ooceraea biroi* | 369 (Oxley et al., 2014) | 493-509 (McKenzie et al., 2016) |
| Florida carpenter ant | *Camponotus floridanus* | 352 (Zhou et al., 2012) | 434-464 (Zube et al., 2008) |
| Big headed leafcutter ant | *Atta cephalotes* | 376 (Engsontia et al., 2015) | 349 (Kelber et al., 2009) |

\*Note that this estimate was generated using sections rather than whole mount brains and is therefore likely an under-estimate of the true number of glomeruli.

**Supplementary Table 2:**
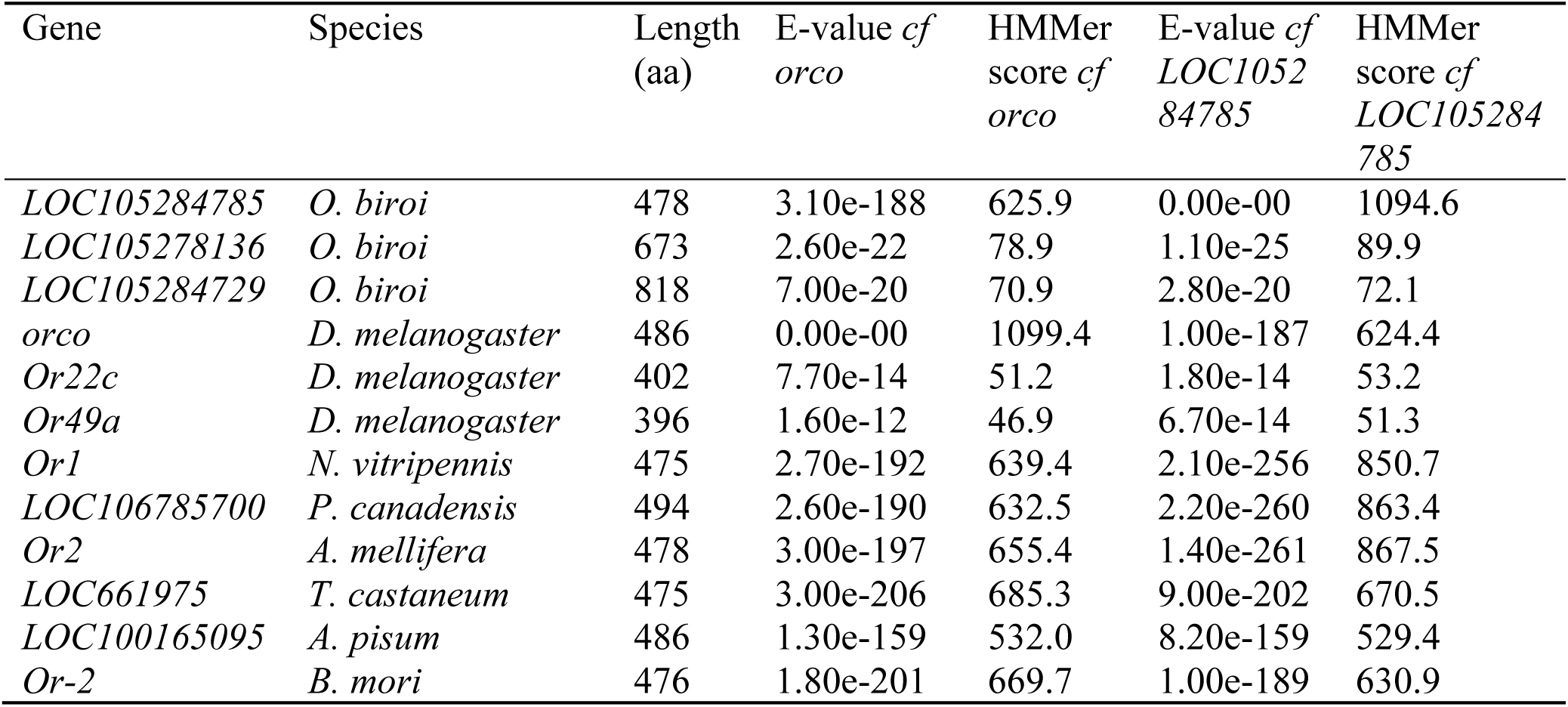
HMMer alignment scores for best matches in eight insect genomes to the *D. melanogaster* (*orco*) and *O. biroi* (*LOC105284785*) *orco* orthologs. The best three matches for *O. biroi* and *D. melanogaster* and the best match for each remaining species are shown. aa: amino acids. Full names of species are: *Ooceraea biroi*, *Drosophila melanogaster*, *Nasonia vitripennis*, *Papilio canadensis*, *Apis mellifera*, *Tribolium castaneum*, *Acyrthosiphon pisum*, and *Bombyx mori*.

| Gene | Species | Length<br>(aa) | E-value *cf*<br>*orco* | HMMer<br>score *cf*<br>*orco* | E-value *cf*<br>*LOC1052*<br>*84785* | HMMer<br>score *cf*<br>*LOC105284*<br>*785* |
| --- | --- | --- | --- | --- | --- | --- |
| *LOC105284785* | *O. biroi* | 478 | 3.10e-188 | 625.9 | 0.00e-00 | 1094.6 |
| *LOC105278136* | *O. biroi* | 673 | 2.60e-22 | 78.9 | 1.10e-25 | 89.9 |
| *LOC105284729* | *O. biroi* | 818 | 7.00e-20 | 70.9 | 2.80e-20 | 72.1 |
| *orco* | *D. melanogaster* | 486 | 0.00e-00 | 1099.4 | 1.00e-187 | 624.4 |
| *Or22c* | *D. melanogaster* | 402 | 7.70e-14 | 51.2 | 1.80e-14 | 53.2 |
| *Or49a* | *D. melanogaster* | 396 | 1.60e-12 | 46.9 | 6.70e-14 | 51.3 |
| *Or1* | *N. vitripennis* | 475 | 2.70e-192 | 639.4 | 2.10e-256 | 850.7 |
| *LOC106785700* | *P. canadensis* | 494 | 2.60e-190 | 632.5 | 2.20e-260 | 863.4 |
| *Or2* | *A. mellifera* | 478 | 3.00e-197 | 655.4 | 1.40e-261 | 867.5 |
| *LOC661975* | *T. castaneum* | 475 | 3.00e-206 | 685.3 | 9.00e-202 | 670.5 |
| *LOC100165095* | *A. pisum* | 486 | 1.30e-159 | 532.0 | 8.20e-159 | 529.4 |
| *Or-2* | *B. mori* | 476 | 1.80e-201 | 669.7 | 1.00e-189 | 630.9 |

Supplementary Table 3: Genotypes and raw data for all ants included in the present study. Supplementary Table 3 is available in the Supplementary Materials. For an explanation of which ants were used in which experiments, see Methods. #-XX (e.g., 3-GO) refers to G1 ants in the datasets from the 5 experimental colonies. # gives the colony number, and XX gives the color tags of the individual ants. These colonies were used for all automated behavioral tracking and survival data (Figures 3D-H, and 4B). Some of these ants were also used for glomerulus counts, antennal lobe volume, and egg-laying data (Figures 2A, 2B, 4A). E-# (e.g., E-1) refers to additional G1 ants used for antennal lobe volume and egg-laying data (Figures 2B and 4A). A-# (e.g., A-1) refers to wild-type ants used for glomerulus counts and antennal lobe volume (Figure 2A and 2B). S-# (e.g. S-1) refers to G2 ants used for Sharpie assays (Figure 3A-C).

**Supplementary Video 1: Antennal lobe reconstructions of wild-type and *orco*^-/-^ adults in *O. biroi* and *D. melanogaster***.

Antennal lobes are reduced in *orco*^-/-^ relative to wild-type in ants, but not in flies. T7 glomeruli in *O. biroi* are labeled in white. Size standards are 20 μm.

**Supplementary Video 2: Sharpie assay.**

Videos of exemplar wild-type and *orco*^-/-^ ants in Sharpie assays (using the same individuals as in Figure 3A,B). The wild-type ant is strongly repelled by Sharpie lines (red) but the *orco*^-/-^ ant is not. Videos are of 2 minute assays playing back at 4x speed.

**Supplementary Video 3: Wandering behavior.**

Automated behavioral tracking of an experimental colony, showing trajectories and identification of ants based on the following color tags: O: orange, P: pink, G: green and B: blue. Note that individual identities are lost when ants form clusters. This two minute video of an experimental colony shows the wandering behavior of *orco*^-/-^ ants (PP and GG; the first letter refers to the thorax tag, the second letter to the gaster tag). OP, which never leaves the nest cluster and therefore is never identified, is *orco*^wt/-^; all other ants are wild-type. PP and GG are seen antennating each other at 0:57 s.

**Supplementary Video 4: Additional social behaviors.**

Ants are the same as in Supplementary Video 3. This video initially shows an *orco*^-/-^ ant, PP, grooming a pile of eggs. At 0:31 s PP begins to move, seemingly eliciting an alarm response that causes the other ants in the colony to move toward PP. Note that GG, the second *orco*^-/-^ ant in the colony, does not become active. PP then moves toward the edge of the dish, possibly depositing a chemical pheromone trail that is followed by some of her nestmates. Note that by the end of the video a clear pheromone trail has formed that many of the wild-type ants are following.

